# Comparative Assessment of *Chlorella vulgaris* Cultivation in Synthetic, Oligotrophic and Industrial Wastewater for Sustainable Biomass Production

**DOI:** 10.1101/2025.07.16.665061

**Authors:** Manu Upadhyay, Vijetna Singh, Neha Singh, Bhumi Nath Tripathi

## Abstract

This study evaluates the physiological and biochemical responses of *Chlorella vulgaris* cultivated in synthetic BG11 medium, oligotrophic water, and industrial wastewater under controlled laboratory conditions with optimized pH and nitrogen-to-phosphorus (N:P) ratios. The highest cell density, growth rate, and pigment accumulation were observed in BG11 at neutral pH and higher N:P ratios. Moderately adjusted N:P ratios in industrial wastewater produced comparable biomass yields and improved photosynthetic efficiency, demonstrating its potential as a cost-effective medium for integrated wastewater treatment and algal biomass production. In contrast, oligotrophic water severely limited growth due to nutrient deficiency. FTIR analyses indicated that *C. vulgaris* maintains essential biochemical components under nutrient stress through adaptive metabolism. This comparative study provides new insights for designing scalable, sustainable microalgal cultivation systems that link biomass production with wastewater management within a circular bioeconomy framework.

## 1. Introduction

Microalgae such as *Chlorella vulgaris* have gained increasing attention as renewable biological resources for addressing environmental and energy challenges, including climate change, fossil fuel depletion, and water pollution (Panahi et al., 2019; Pittman et al., 2011). *C. vulgaris* is widely studied due to its high growth rate, adaptability, and ability to accumulate valuable compounds such as proteins, lipids, and pigments (Ratha et al., 2012). Its capacity to thrive in nutrient-poor and nutrient-rich waters makes it a promising candidate for combined biofuel production and wastewater treatment (Umamaheswari et al., 2016).

Synthetic media such as BG11 are well-established for laboratory cultivation of microalgae, providing controlled nutrient conditions and reproducible growth (Rippka et al., 1979; Wang et al., 2023). However, the cost and sustainability of large-scale cultivation remain key constraints. Using alternative water sources, such as oligotrophic water or industrial wastewater, could reduce costs and enable resource recovery, but these media often have imbalanced or low nutrient levels (Rawat et al., 2011; Wang et al., 2010; Yang et al., 2022). Although previous studies have demonstrated the feasibility of growing *C. vulgaris* in wastewater, few have systematically compared its growth and biochemical responses in synthetic, natural, and industrial media under identical, well-controlled conditions. In particular, the combined effects of pH and nitrogen-to-phosphorus (N:P) ratios on growth performance and metabolic adjustment remain underexplored (Pandey et al., 2023; Kundu et al., 2024).

Maintaining optimal pH is critical for nutrient solubility, enzyme activity, and photosynthesis (Kim et al., 2014; Wang et al., 2016). Similarly, adjusting the N:P ratio can enhance algal productivity, especially in wastewater with variable nutrient compositions (Xin et al., 2010; Zhou et al., 2012). Yet, evidence from rigorously controlled side-by-side comparisons is limited.

This study addresses this gap by comparing *C. vulgaris* cultivation in BG11, oligotrophic water, and industrial wastewater while systematically adjusting pH and N:P ratios under controlled laboratory conditions. By analyzing growth kinetics, photosynthetic efficiency, pigment accumulation, protein content, and biochemical composition, this work provides reproducible data to guide sustainable algal biomass production and integrated wastewater management within a circular bioeconomy framework.

## 2. Material Methods

### 2.1 Microalgal Strain, Culture Media, Experimental Design

The green microalga *Chlorella vulgaris* was selected for this study due to its robust growth and adaptability across diverse culture media (Ratha et al., 2012). Three types of media were used: BG11 (a standard synthetic algal medium), oligotrophic water, and industrial wastewater. All media were prepared and sterilized according to established protocols (Stanier et al., 1971; Wang et al., 2010). Cultures were initially inoculated in each medium and maintained under standard light and temperature conditions as described by Tripathi et al. (2004). Sampling and analyses were performed at 0, 3, 6, 9, and 12 days to monitor acclimation and growth dynamics. The cultures were grown under controlled laboratory conditions with systematic variations in pH and nitrogen-to-phosphorus (N:P) ratios to evaluate their effects on growth and physiological responses. pH and N:P ratios were adjusted using analytical-grade reagents and monitored throughout the experiment (Xin et al., 2010). All cultures were inoculated at an initial cell density of 1 × 10⁴ cells mL⁻¹ and incubated in 250 mL Erlenmeyer flasks containing 75 mL of the respective medium.

#### 1. Cell Count Determination

Cell density was measured 0,3,6,9 and 12 days using a Neubauer haemocytometer under a light microscope. The cell count was calculated using the formula:

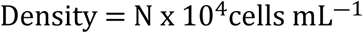

where N is the number of cells counted (Padang et al, 2018)

### 2.2 Growth Rate Estimation

Growth rates were determined spectrophotometrically by measuring the optical density at 663 nm Thermo Scientific Orion AquaMate 8100 UV Visible spectrophotometer. The specific growth rate (μ) was calculated as:

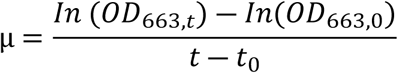

where OD_663,0_ and OD_663,t_ are the optical densities at the initial and final time points, respectively (Haslianti et al, 2023; Richmond et al, 2004).

### 2.3 Determination of Biomass and Biomass Yield

The biomass of the selected algal species was collected after 12 days of inoculation. The samples were centrifuged using a Hermle Z366 centrifuge at 4100X*g* for 10 min at room temperature. The pellets obtained by centrifugation were washed with distilled water and dried. Biomass yield was expressed as % increase per day per liter, following standard methods (Markou et al, 2013).

The biomass percentage was determined using the following equation:

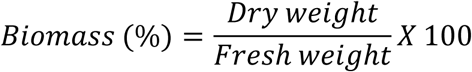

To determine the biomass yield, the following equation was applied (Shalaby et al 2011):

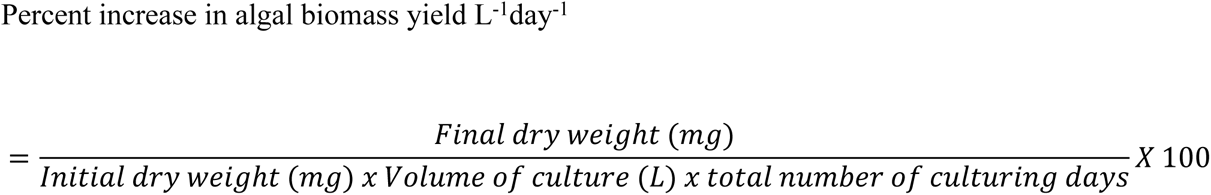

### 2.4 Photosynthetic Efficiency (Fv/Fm)

The maximum quantum yield of photosystem II (Fv/Fm) was measured using a pulse-amplitude modulated fluorometer MINI-PAM-II (Photosynthesis Yield Analyzer), WALZ, after dark adaptation for 15 minutes, as described by Maxwell et al. (2000).

### 2.5 Quantification of Photosynthetic Pigments

Chlorophyll a and carotenoids were extracted from algal samples following the method of Arnon (1949). A 5 mL aliquot of algal culture was harvested and centrifuged at 5000 rpm for 10 minutes at room temperature using a Hermle Z366 centrifuge. The resulting pellet was resuspended in 80% acetone and stored at 4°C for 24 hours. After storage, the sample was centrifuged again, and the absorbance of the supernatant was measured at 663, 645, and 480 nm using a Thermo Scientific Orion AquaMate 8100 UV–Visible spectrophotometer. Chlorophyll a and carotenoid contents were estimated using the following equation:

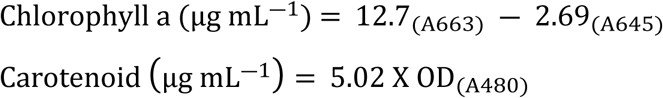

### 2.6 Determination of Protein

The protein content was quantitatively determined following the method of Lowry et al. (1951), with absorbance readings taken using a Thermo Scientific Orion AquaMate 8100 UV–visible spectrophotometer.

## 3. Results

### 3.1 Growth in terms of cell count with variations in pH and N:P ratio

The cell count analysis of *Chlorella vulgaris* over the 12-day incubation period revealed distinct growth patterns influenced by both pH and N:P ratios (Figure 1). In BG11 medium (Figure 1a), the highest cell density was recorded at pH 7, while acidic (pH 5) and alkaline (pH 9) conditions resulted in slightly lower yields, indicating that near-neutral pH supports optimal growth in nutrient-rich conditions. In oligotrophic water (Figure 1b) and industrial wastewater (Figure 1c), overall cell counts were lower than in BG11, but pH 7 again supported relatively higher densities, suggesting that neutral pH promotes better adaptation even under nutrient limitations. Regarding the N:P ratio, BG11 medium (Figure 1d) showed a clear increase in cell count with higher N:P ratios, with the 30:1 ratio achieving the highest yield by day 12, suggesting that additional nitrogen can further boost biomass production in an already nutrient-rich medium. Similarly, in oligotrophic water (Figure 1e) and wastewater (Figure 1f), the highest N:P ratio (30:1) resulted in slightly increased cell counts compared to the control and lower ratios.

**Figure 1.**
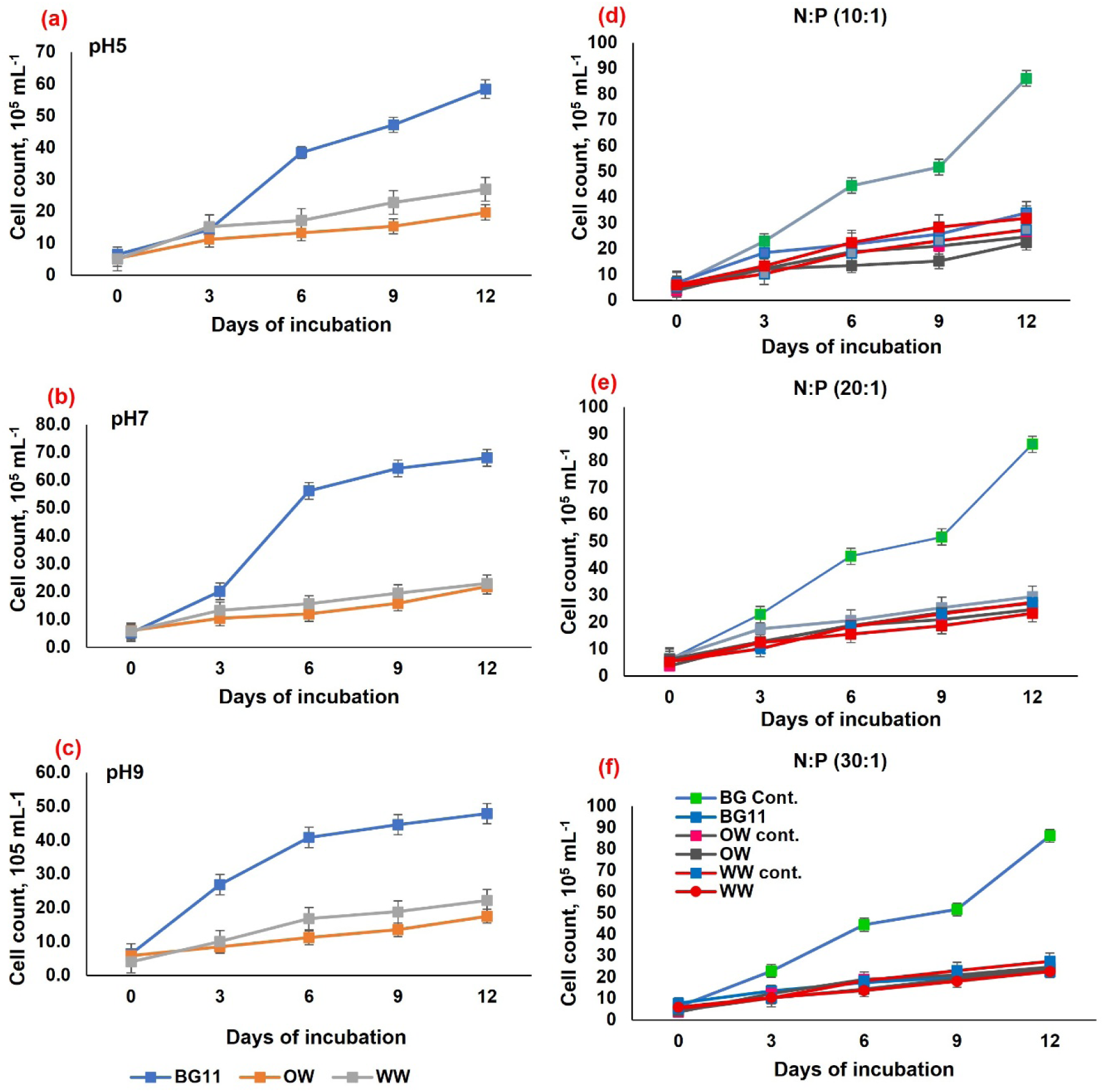
Cell count (10^4^ cells mL^-1^) of *Chlorella vulgaris* grown in different media with variations of pH and N:P ratio

### 3.2 Specific Growth Rate of *Chlorella vulgaris* in Oligotrophic Water (OW) and Wastewater (WW) Media

The specific growth rate (μ) of *Chlorella vulgaris* showed distinct patterns in oligotrophic water (OW) and industrial wastewater (WW) compared to the conventional BG11 medium across different pH conditions (Figure 2a-c). In the nutrient-rich BG11 medium (Figure 2a), the control condition-representing the standard near-neutral pH-supported the highest and most sustained growth rate, while shifting the pH to acidic (pH 5) or alkaline (pH 9) levels caused a marked decline in growth after day 6. Similarly, in OW (Figure 2b) and WW (Figure 2c), the highest specific growth rates were recorded at pH 7, whereas both acidic and especially alkaline conditions resulted in reduced growth. However, the overall growth differed between the two natural waters: in WW, growth rates were substantially higher than in OW, particularly at near-neutral pH, and showed rapid adaptation with a notable increase around day 6-7. In contrast, the limited nutrient availability in OW led to consistently lower specific growth rates across all pH levels, with a prolonged lag phase and a less distinct exponential phase, especially under acidic and alkaline conditions.

**Figure 2.**
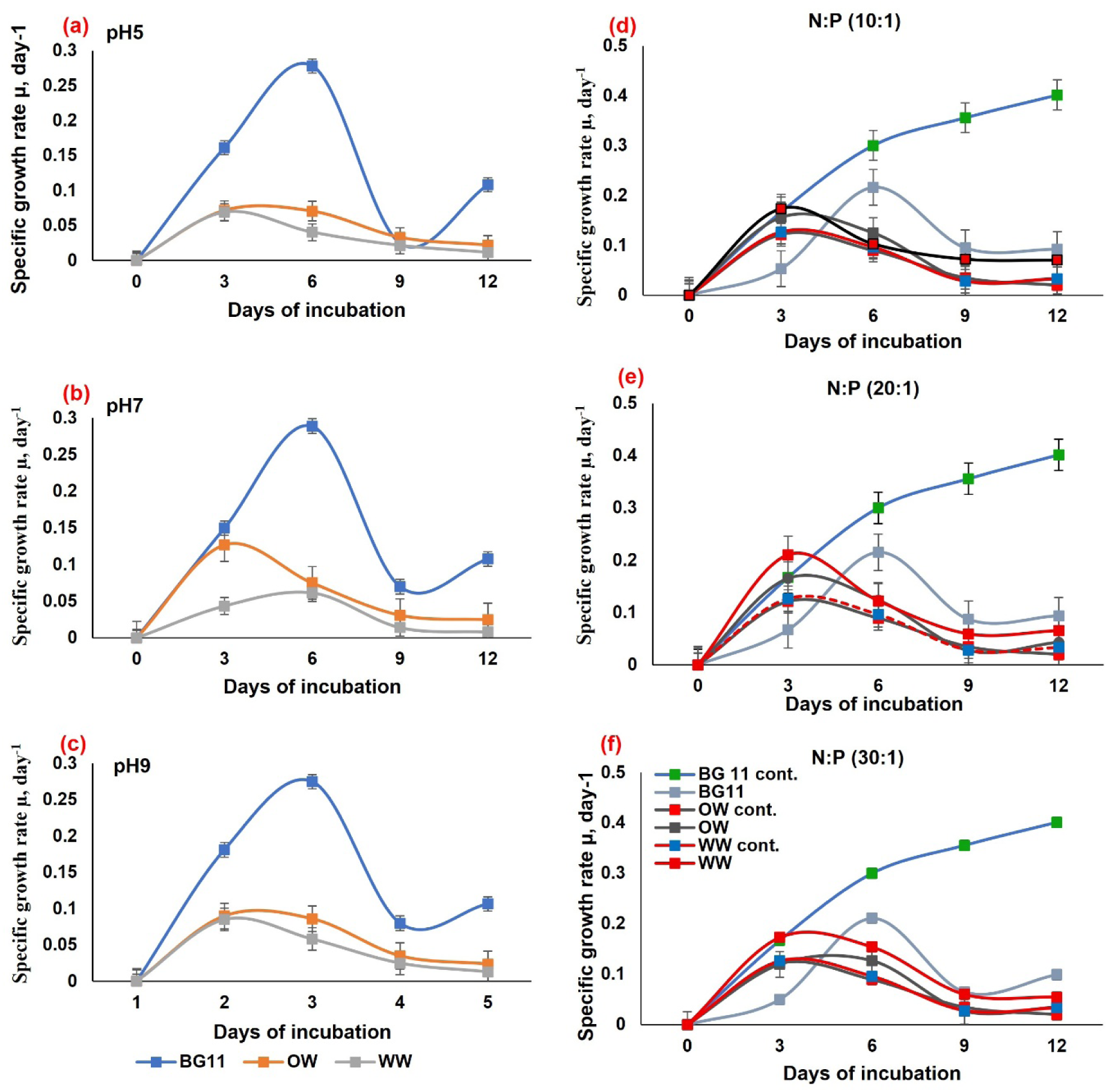
Specific growth rate (μ) of *Chlorella vulgaris* grown in different media with variations of pH and N:P ratio

Figure 2 (d–f) illustrates how varying the nitrogen-to-phosphorus (N:P) ratio and growth medium affects the specific growth rate of *Chlorella vulgaris*. In the nutrient-rich BG11 medium (Figure 2d), the control condition showed the highest and most sustained growth rate, while increasing the N:P ratio to 10:1, 20:1, or 30:1 slightly reduced growth. This indicates that the standard BG11 formulation already provides an optimal nutrient balance. In contrast, in oligotrophic water (Figure 2e) and wastewater (Figure 2f), where nutrients are more limited or imbalanced, adjusting the N:P ratio had a clear positive effect. A moderate N:P ratio of 20:1 produced the highest growth rate in both OW and WW, outperforming the control and other N:P treatments.

### 3.3 Biomass Percentage and Biomass Yield Percentage

The biomass percentage and yield results (Figure 3) clearly demonstrate that both the nutrient status of the medium and variations in pH or N:P ratio significantly affect *Chlorella vulgaris* productivity. In the nutrient-rich BG11 medium, the highest biomass percentage and daily yield were consistently recorded at pH 7 and under the control condition without additional nutrient adjustments, confirming that a balanced nutrient supply and neutral pH maximize growth under ideal conditions. When the same treatments were applied in oligotrophic water (OW) and wastewater (WW), both biomass percentage and yield declined noticeably compared to BG11. OW, being nutrient-poor, supported the lowest biomass and yield across all treatments, indicating that limited baseline nutrients strongly restrict productivity regardless of pH or N:P ratio. In contrast, WW produced slightly higher biomass and yield than OW, demonstrating that it supplies more nutrients but still less than BG11. Notably, in WW, adjusting the N:P ratio to a moderate level (20:1) increased both biomass percentage and yield compared to the control, highlighting that targeted nutrient enrichment can partly offset nutrient imbalances in wastewater and enhance microalgal productivity.

**Figure 3.**
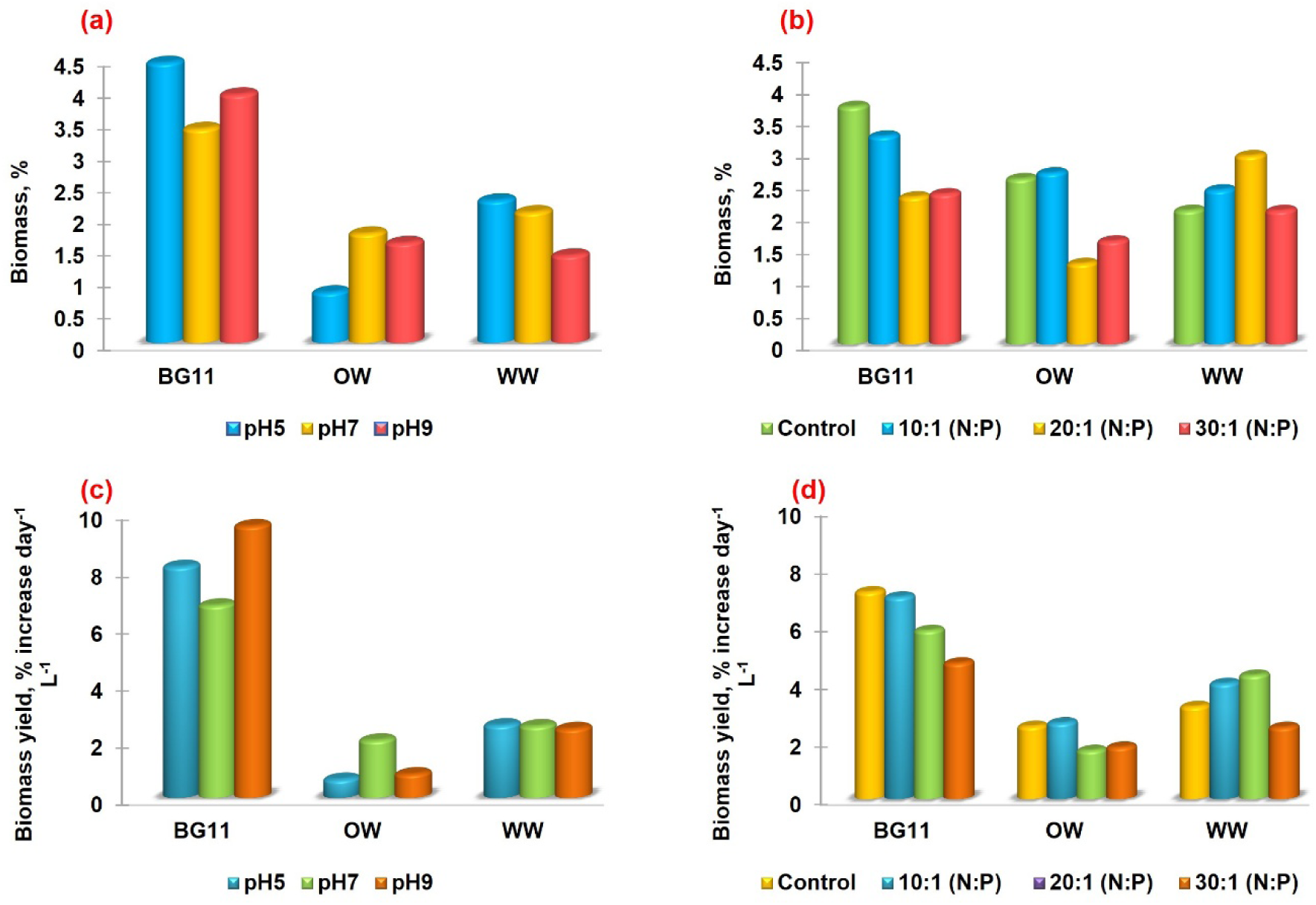
Biomass % and Biomass yield % of *Chlorella vulgaris* grown in different media with variations of pH and N:P ratio

### 3.4 Fv/Fm (Maximum Quantum Yield of Photosystem II)

The maximum quantum yield of photosystem II (Fv/Fm) in *Chlorella vulgaris* was strongly influenced by pH, nutrient medium composition, and N:P ratio variations (Figure 4). In BG11 medium, Fv/Fm increased rapidly across all pH and N:P treatments, remaining above 0.70, which indicates healthy PSII performance. Mildly alkaline conditions (pH 9) slightly enhanced Fv/Fm further. In OW and WW, overall Fv/Fm values were lower than in BG11, reflecting nutrient limitations that constrain photosynthetic efficiency. However, in both OW and WW, maintaining a neutral pH or the control condition resulted in slightly higher Fv/Fm values than acidic or alkaline conditions (Figure 4b, c). For N:P treatments, moderate enrichment (10:1 or 20:1) produced clear peaks in Fv/Fm in OW and WW (Figure 4e, f), indicating that balanced nitrogen and phosphorus levels can temporarily improve photosynthetic efficiency under nutrient-limited conditions. By contrast, the highest N:P ratio (30:1) did not provide additional benefit and in some cases reduced Fv/Fm, likely due to nutrient imbalance.

**Figure 4.**
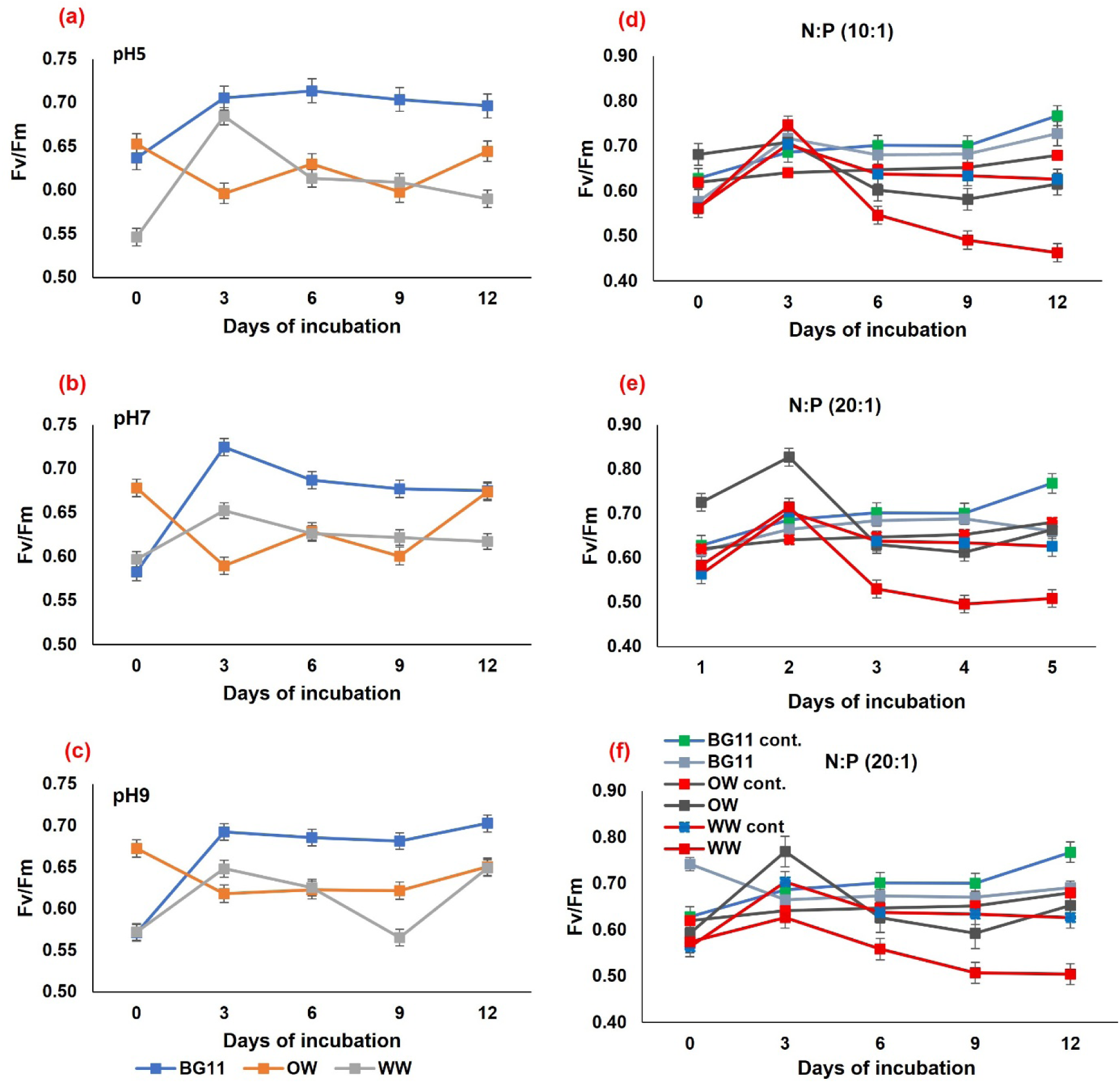
Fv/Fm of *Chlorella vulgaris* grown in different media with variations of pH and N:P ratio

### 3.5 Pigment Content

The pigment analysis (Figure 5) shows how pH and N:P ratio affect carotenoid and chlorophyll *a* content in *Chlorella vulgaris*. In BG11, carotenoid content peaked at pH 5 (Figure 5a) and with moderate N:P enrichment (10:1) (Figure 5b), indicating that mild stress or balanced nitrogen can boost protective pigment accumulation. Chlorophyll *a* content was highest under alkaline conditions (pH 9) and at an N:P ratio of 10:1 (Figure 5c, d), suggesting that slightly higher pH and nitrogen availability enhance photosynthetic pigment synthesis in nutrient-rich media. In contrast, OW showed consistently low carotenoid levels regardless of pH or N:P ratio, but chlorophyll *a* increased significantly at pH 9 and with moderate N:P enrichment (10:1), demonstrating that even a nutrient-poor medium can partially support pigment formation when conditions are optimized. However, WW consistently showed very low carotenoid and chlorophyll *a* levels under all treatments, indicating that the nutrient balance and quality of the wastewater used in this study were insufficient to support significant pigment accumulation.

**Figure 5.**
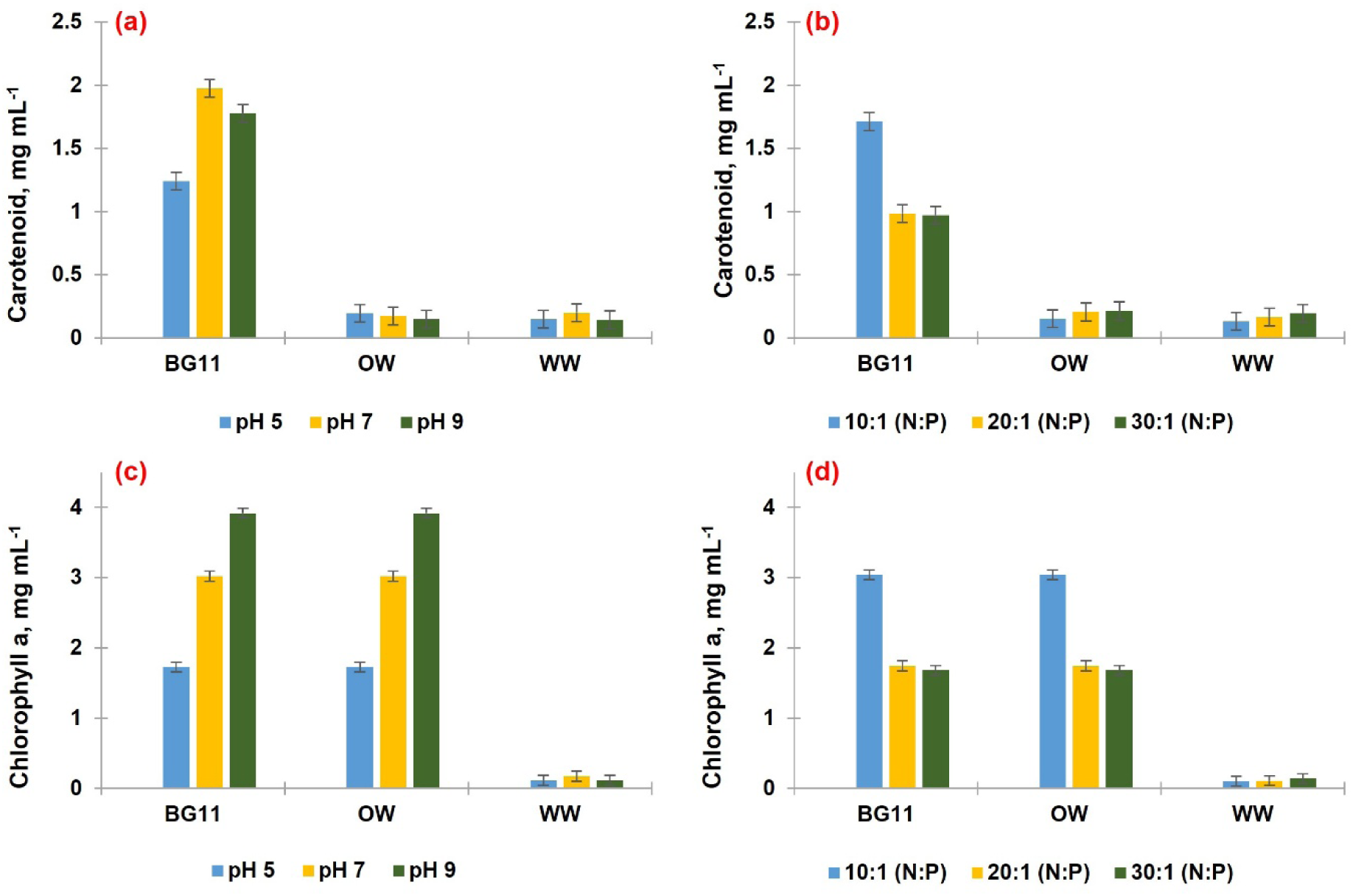
Chlorophyll a and carotenoid content (mg L^-1^) of *Chlorella vulgaris* cultivated in different media under varying pH levels and N:P ratios.

### 3.6 Protein Content

The protein content results (Figure 6) indicate that neither pH adjustment nor N:P ratio variation had a significant effect on total protein accumulation in *Chlorella vulgaris* across the different media. In BG11, protein levels remained stable at around 1.0 mg L⁻¹ under all pH conditions, with only minor fluctuations observed at pH 5 or pH 9 (Figure 6a). Similarly, in OW and WW, protein concentrations were consistently lower than in BG11 (approximately 0.8–0.9 mg L⁻¹) and showed minimal variation between pH treatments, suggesting that protein synthesis remains relatively stable despite pH stress in nutrient-limited environments. For the N:P ratio treatments (Figure 6b), protein content in BG11 showed a slight decline from the control (∼1.0 mg L⁻¹) with increasing N:P ratios (10:1, 20:1, 30:1), but this decrease was minimal. In OW and WW, protein concentrations again remained largely unchanged across all N:P ratios, indicating that additional nitrogen and phosphorus did not substantially increase protein content under low-nutrient or wastewater conditions.

**Figure 6.**
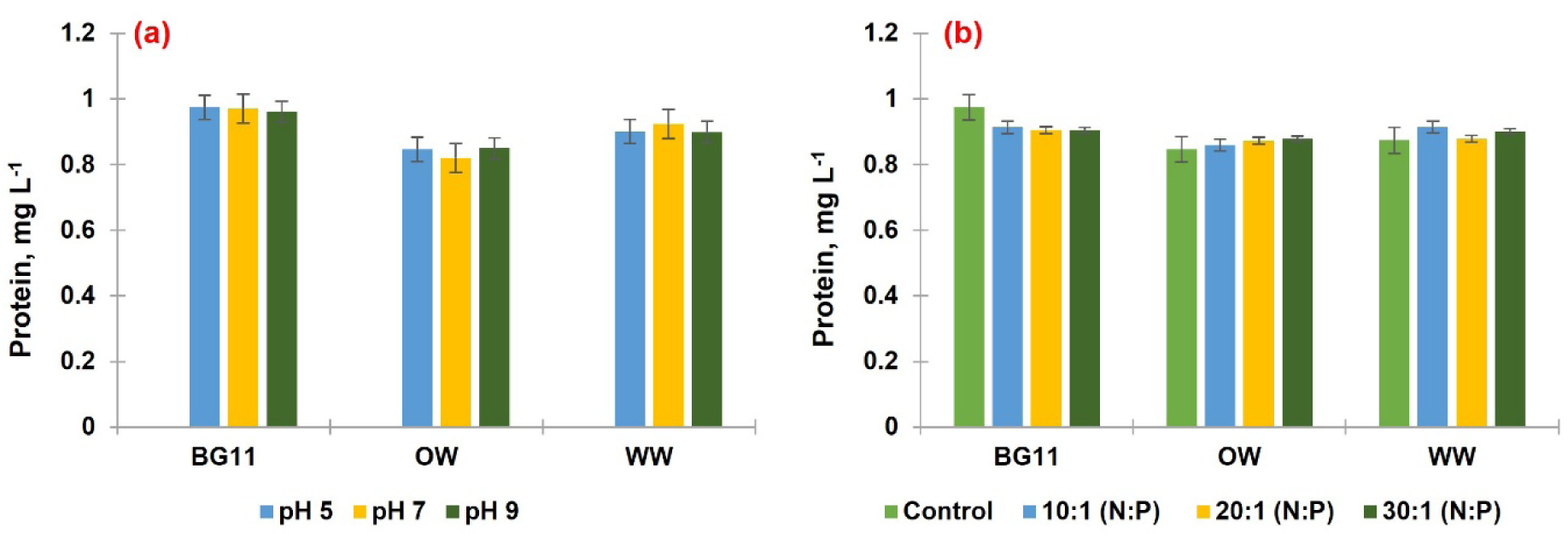
Protein (mg mL^-1^) of *Chlorella vulgaris* grown in different media with variations of pH and N:P ratio

### 3.7 FTIR analysis

The FTIR spectra (Figures 7–9) show that *Chlorella vulgaris* cultivated in BG11, oligotrophic water (OW), and wastewater (WW) displayed distinct yet overlapping functional groups, reflecting variations in biochemical composition under different pH and N:P ratio treatments. Across all media, key peaks were consistently observed in regions associated with water and proteins (3270–3371 cm⁻¹), lipids (around 2921–2922 cm⁻¹), aldehydes and alkenes (2851– 2854 cm⁻¹), and proteins (amide I and II bands near 1630–1647 cm⁻¹ and ∼1540 cm⁻¹). Compared to BG11, OW and WW showed slight shifts in peak positions but retained the main functional groups, indicating that nutrient-poor or wastewater conditions did not eliminate major biochemical features but did influence macromolecular structure. Notably, peaks related to nucleic acids (1372–1398 cm⁻¹), phosphate groups (1240–1250 cm⁻¹), and polysaccharides (around 1015–1045 cm⁻¹) were present across all treatments but showed minor shifts in OW and WW, suggesting changes in cell wall or nucleic acid content under nutrient stress. Small differences were also seen for nitrite peaks (∼2360 cm⁻¹) and alkyl or aromatic amines (1148– 1151 cm⁻¹), indicating subtle biochemical adjustments. The presence of unsaturated aldehydes and ketones (around 1740–1743 cm⁻¹) in all media confirms active lipid and carbonyl metabolism, while variations in peak intensities and slight shifts suggest that OW and WW conditions modestly modified the biochemical profile, likely due to nutrient limitations or stress adaptation.

**Figure 7.**
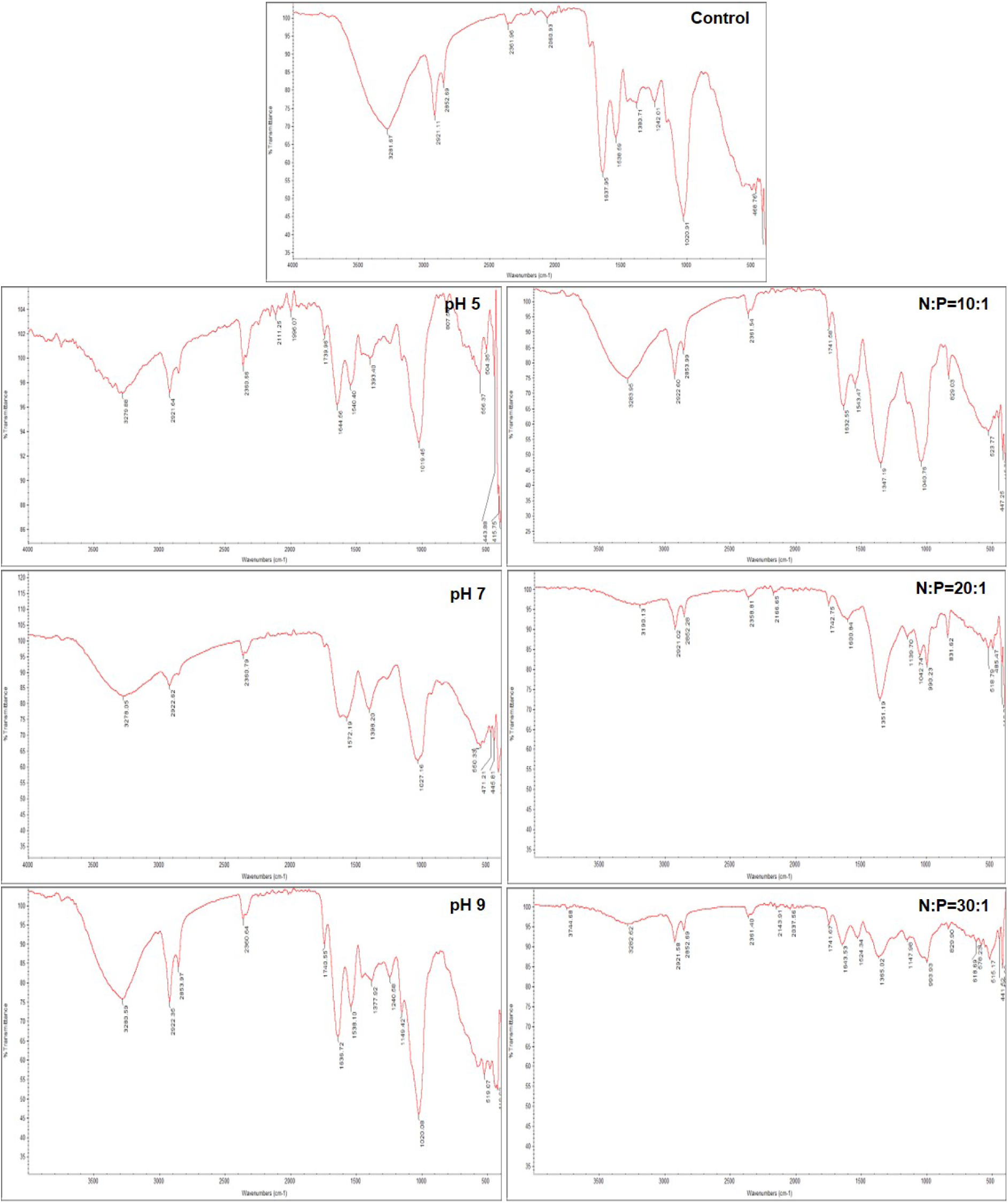
FTIR Spectra of *Chlorella vulgaris* grown in BG11 media with different variations of pH and N:P ratio

**Figure 8.**
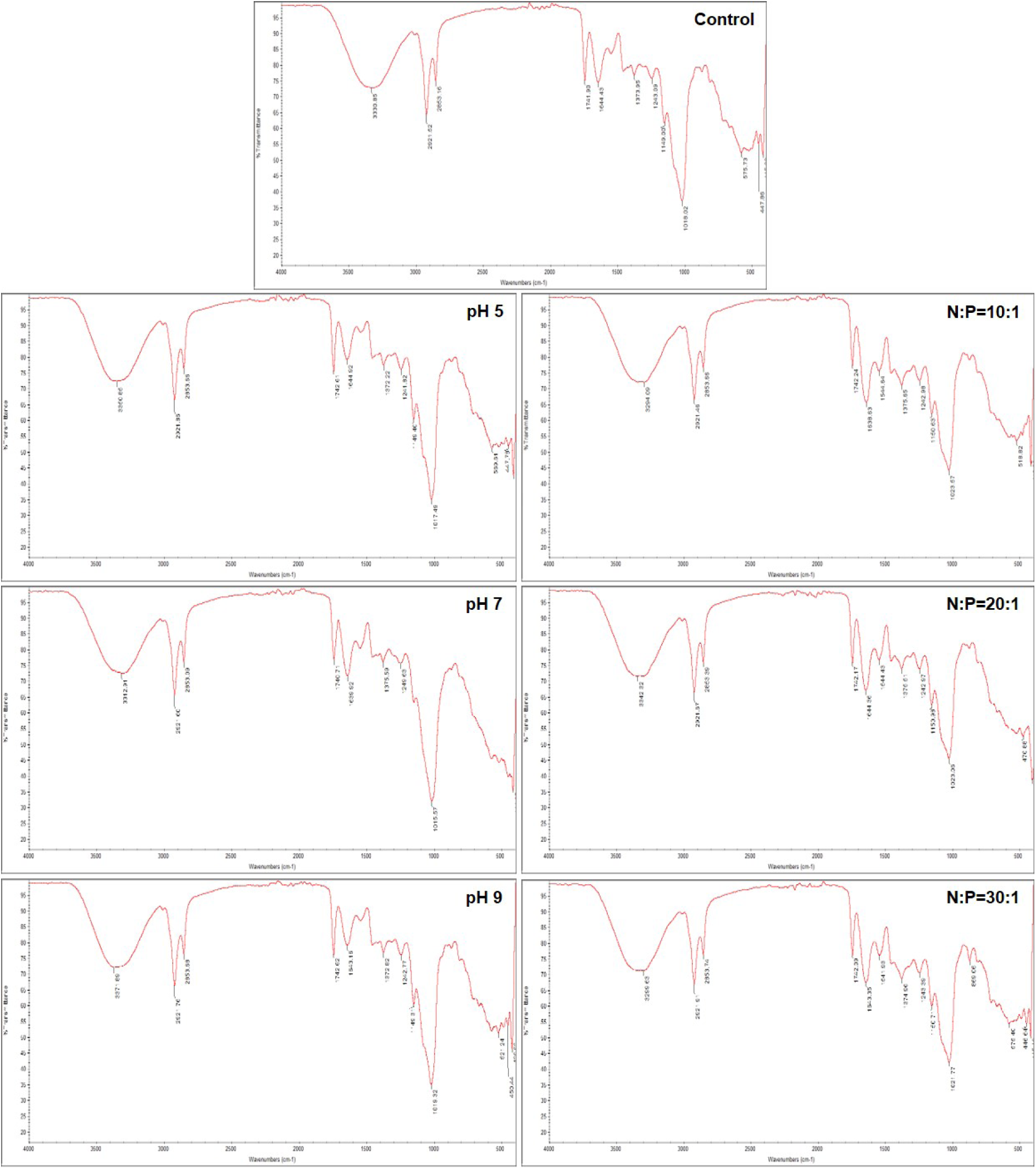
FTIR Spectra of *Chlorella vulgaris* grown in Oligotrophic water with different variations of pH and N:P ratio

**Figure 9.**
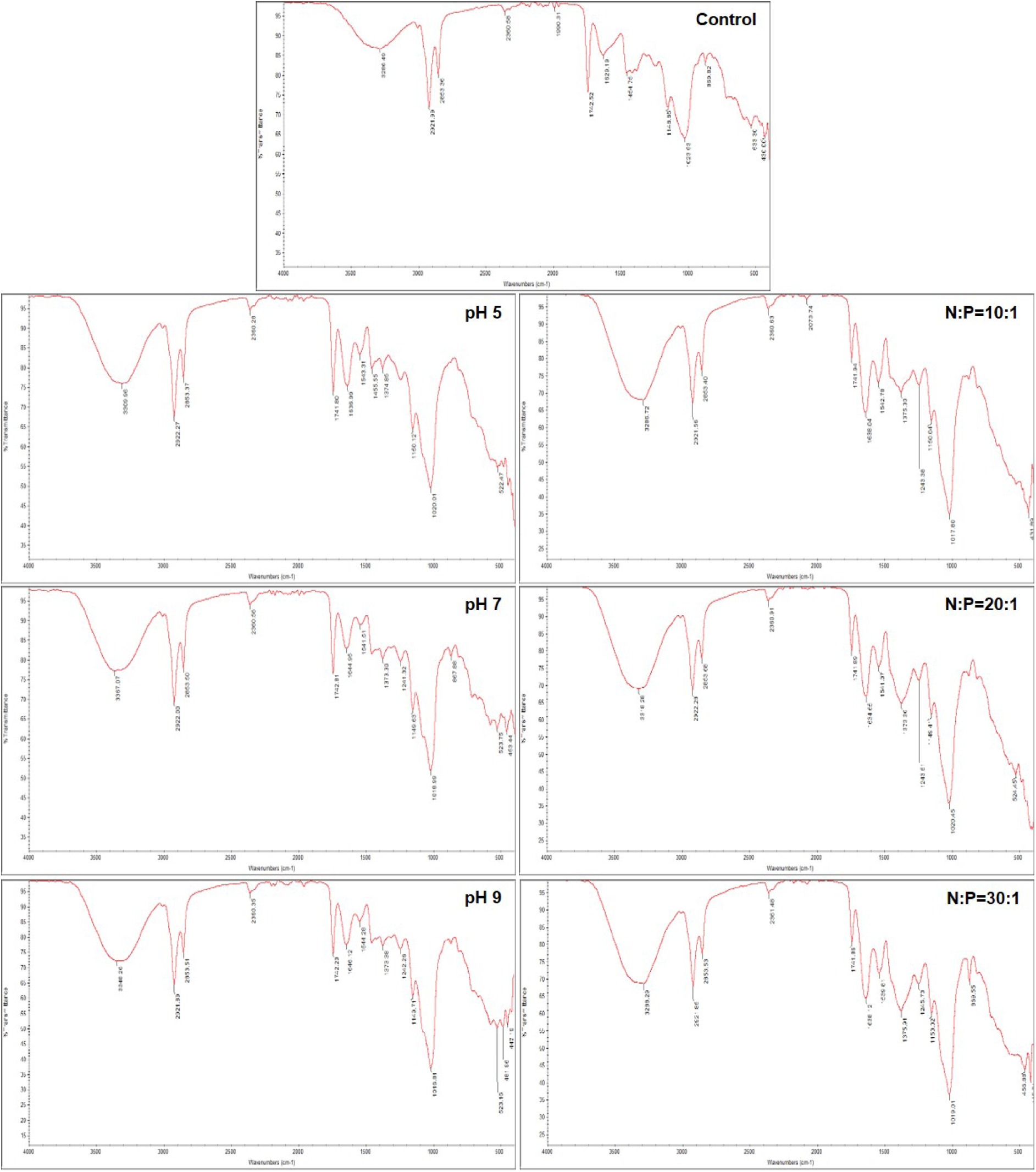
FTIR Spectra of *Chlorella vulgaris* grown in Waste water with different variations of pH and N:P ratio

## Discussion

This study confirms that both pH and N:P ratio are critical factors influencing the growth and physiological responses of *Chlorella vulgaris* cultivated in synthetic, oligotrophic, and wastewater media. In BG11, the highest cell densities and specific growth rates at neutral pH align with earlier findings that near-neutral pH optimizes enzyme activity, nutrient availability, and photosynthetic performance (Wang et al., 2016). The decline in growth under acidic or alkaline conditions supports the well-known sensitivity of microalgae to pH stress (Kim et al., 2014; Solovchenko et al., 2019).

In comparison, both oligotrophic water and wastewater supported lower biomass yields than BG11, highlighting the importance of adequate nutrient supply for sustaining microalgal productivity (Zhou et al., 2011). However, the observation that neutral pH still promoted relatively higher growth even in nutrient-limited conditions suggests that pH control can partially compensate for nutrient deficiencies by improving nutrient solubility and metabolic activity (Yu et al., 2022).

The effects of the N:P ratio further demonstrate how nutrient stoichiometry shapes algal growth. In BG11, the standard formulation appeared sufficient, as higher N:P ratios did not improve growth significantly. Conversely, moderate N:P enrichment (20:1) in wastewater enhanced both cell counts and biomass yield, indicating that targeted nutrient balancing can partially overcome nutrient imbalances typical of industrial effluents (Xin et al., 2010; Zhou et al., 2012). This finding supports the practical use of wastewater as a low-cost growth medium when combined with simple nutrient supplementation, contributing to integrated wastewater treatment and biomass recovery (Pittman et al., 2011).

Photosynthetic efficiency trends, as indicated by Fv/Fm values, reinforce the impact of both medium composition and environmental parameters. High Fv/Fm values in BG11 confirm healthy PSII activity under balanced conditions (Maxwell & Johnson, 2000), while lower values in OW and WW reflect nutrient constraints that limit photochemical capacity (Yee et al., 2025). The temporary improvements in Fv/Fm at moderate N:P ratios in OW and WW demonstrate that balanced nutrient availability can help maintain photosynthetic performance, although excessive enrichment (30:1) may cause metabolic imbalance (Geider & La Roche, 2002).

The pigment profiles show that mild stress or moderate nitrogen enrichment can stimulate protective carotenoid production and chlorophyll *a* accumulation, but only when baseline nutrients are sufficient (Solovchenko et al., 2013; Panahi et al., 2019). In wastewater, the consistently low pigment levels suggest that its nutrient composition alone was inadequate to support robust pigment biosynthesis, underlining the need for targeted nutrient optimization. Interestingly, total protein content remained relatively stable across all treatments, regardless of pH or N:P ratio. This suggests that protein synthesis in *C. vulgaris* is less sensitive to moderate environmental changes than growth or pigment production, reflecting metabolic regulation under stress (Markou & Nerantzis, 2013).

The FTIR results further confirm that while the main functional groups were conserved across treatments, minor peak shifts in OW and WW indicate subtle biochemical adjustments and stress responses. The persistence of protein, lipid, polysaccharide, and nucleic acid signatures under different conditions highlights the metabolic plasticity of *C. vulgaris*, making it well-suited for use in variable or lower-cost water sources when properly managed (Pandey et al., 2010).

Overall, this study fills an important knowledge gap by providing reproducible, side-by-side evidence of how *C. vulgaris* responds physiologically and biochemically to contrasting water sources under systematically controlled pH and nutrient conditions. These findings strengthen the basis for developing scalable, circular bioeconomy systems that couple wastewater treatment with cost-effective algal biomass production.

## Conclusion

This study demonstrates that pH and N:P ratio are key factors shaping the growth and biochemical responses of *Chlorella vulgaris* across different culture media. Neutral pH and balanced nutrient conditions consistently supported higher biomass yields, photosynthetic efficiency, and pigment production in BG11 and, to a lesser extent, in wastewater. The results confirm that industrial wastewater, when combined with moderate nutrient adjustment, can serve as a viable low-cost medium for sustainable algal cultivation and integrated wastewater treatment. The comparative approach and systematic controls presented here provide practical insights for developing scalable microalgal systems within a circular bioeconomy framework.

## Acknowledgements

Financial support from the DBT, New Delhi in the form of a DBT-BUILDER-IGNTU project (Grant No. BT/INF/22/SP45361/2022) is gratefully acknowledged. We are also thankful to DST, New Delhi for financial assistance for renovation of Algal Culture Room through DST-FIST Project (Grant No. SR/FST/LS-1/2021/888).

## Data Availability

All data generated or analyzed during this study are included in this published article and its supplementary information files.

## Declarations

## Ethical Approval and Consent to Participate

Not applicable.

